# Diffusion Histology Imaging Combining Diffusion Basis Spectrum Imaging (DBSI) and Machine Learning Improves Detection and Classification of Glioblastoma Pathology

**DOI:** 10.1101/843367

**Authors:** Zezhong Ye, Richard L. Price, Xiran Liu, Joshua Lin, Qingsong Yang, Peng Sun, Anthony T. Wu, Liang Wang, Rowland Han, Chunyu Song, Ruimeng Yang, Sam E. Gary, Diane D. Mao, Michael Wallendorf, Jian L. Campian, Jr-Shin Li, Sonika Dahiya, Albert H. Kim, Sheng-Kwei Song

**Author notes:** Corresponding Authors: Albert H. Kim, M.D., Ph.D., Department of Neurological Surgery, Washington University School of Medicine, Campus Box 8057, 660 S. Euclid Avenue, St. Louis, MO 63110, USA, Email address, Sonika Dahiya, M.D., Department of Pathology and Immunology, Washington University School of Medicine, 660 S. Euclid Avenue, St. Louis, MO 63110, USA. Institute of Computational and Mathematical Engineering, Stanford University, Palo Alto, CA 94305, USA. Keck School of Medicine, University of Southern California, Los Angeles, CA 90033, USA.

## Abstract

**Purpose:** Glioblastoma (GBM) is one of the deadliest cancers with no cure. While conventional MRI has been widely adopted for examining GBM clinically, accurate neuroimaging assessment of tumor histopathology for improved diagnosis, surgical planning, and treatment evaluation, remains an unmet need in the clinical management of GBMs.

**Experimental Design:** We employ a novel Diffusion Histology Imaging (DHI) approach, combining diffusion basis spectrum imaging (DBSI) and machine learning, to detect, differentiate, and quantify areas of high cellularity, tumor necrosis, and tumor infiltration in GBM.

**Results:** Gd-enhanced T1W or hyper-intense FLAIR failed to reflect the morphological complexity underlying tumor in GBM patients. Contrary to the conventional wisdom that apparent diffusion coefficient (ADC) negatively correlates with increased tumor cellularity, we demonstrate disagreement between ADC and histologically confirmed tumor cellularity in glioblastoma specimens, whereas DBSI-derived restricted isotropic diffusion fraction positively correlated with tumor cellularity in the same specimens. By incorporating DBSI metrics as classifiers for a supervised machine learning algorithm, we accurately predicted high tumor cellularity, tumor necrosis, and tumor infiltration with 87.5%, 89.0% and 93.4% accuracy, respectively.

**Conclusion:** Our results suggest that DHI could serve as a favorable alternative to current neuroimaging techniques for guiding biopsy or surgery as well as monitoring therapeutic response in the treatment of glioblastoma.

**Translational Relevance:** Current clinical diagnosis, surgical planning, and assessment of treatment response for GBM patients relies heavily on gadolinium-enhanced T1-weighted MRI, which is non-specific for tumor growth and merely reflects a disrupted blood-brain barrier. The complex tumor microenvironment and spatial heterogeneity make GBM difficult to characterize using current clinical imaging modalities. In this study, we developed a novel imaging technique to characterize and accurately predict key histological features of GBM - high tumor cellularity, tumor necrosis, and tumor infiltration. While further validation in a larger cohort of patients is needed, the current proof-of-concept approach could provide a solution to resolve important clinical questions such as the identification of true tumor progression vs. pseudoprogression or radiation necrosis.

## Introduction

Glioblastoma (GBM) is the most common primary malignant brain tumor in adults (1). It is estimated that 13,140 new GBM cases will be diagnosed during 2020 in the U.S. (1). Despite extensive multimodality treatment, which includes surgical resection, chemotherapy, and radiation, patients with GBM exhibit a dismal 5-year survival rate of 6.8% (1). Histologically, GBMs are characterized by increased cellularity, vascular proliferation, necrosis, and infiltration into normal brain parenchyma (2). Currently, the histopathological complexity of GBM cannot be fully appreciated without microscopic examination of tumor specimens.

Gadolinium (Gd)-enhanced T1-weighted (T1W) MRI is the standard clinical imaging modality for detection, surgical planning, and evaluation of GBM treatment response (3-6). Contrast-enhancement in T1W images is clinically interpreted as a measure of tumor burden and is widely used as the target for surgical resection (6). However, due to the infiltrative nature of GBM, tumor cells are known to extend well beyond the area of contrast enhancement (3). After treatment, contrast enhancement is not diagnostically specific for GBM, since it reflects not only increased Gd leakage due to angiogenesis induced by malignant tumors but also blood-brain barrier disruption triggered by other factors, including radiation effects and ischemia (7-9). Therefore, Gd enhancement neither accurately measure tumor burden nor specifically reflect various pathological changes.

Conventional MR sequences such as T2-weighted imaging (T2W) and fluid-attenuated inversion recovery (FLAIR) imaging have also been employed to localize non-enhancing tumor to complement Gd-enhanced T1W images. The combination of these imaging sequences was adopted into the Response Assessment in Neuro-Oncology (RANO) (3). However, precisely quantifying the increase in T2W/FLAIR image signal intensity remains difficult. Differentiating non-enhancing tumor from other causes of increased T2W or FLAIR image signal intensity, such as edema, radiation effect, ischemic injury, postoperative changes, or other treatment effects, continues to challenge clinicians.

In addition to conventional T1W and T2W imaging, diffusion-weighted imaging (DWI) and the derived apparent diffusion coefficient (ADC) have been employed to detect and assess tumor cellularity in many cancers based on the hypothesis that increased tumor cellularity restricts diffusion, leading to decreased ADC values.(10) ADC has been shown to decrease with increasing glioma grade (11) and applied to characterize the infiltrative pattern of recurrent tumor after treatment (12). However, ADC loses specificity and sensitivity in the presence of co-existing necrosis (increased ADC), tumor infiltration (decreased ADC) and/or vasogenic edema (increased ADC), complicating local brain diffusion characteristics. Combining multiple MR sequences falls short of predicting the complex and heterogeneous GBM tumor microenvironment. Additionally, the gold standard of surgical biopsy carries risk. Thus, the development of noninvasive alternatives to decipher the complex GBM tumor histopathology is urgently needed so that clinicians can make rational decisions about continuing, stopping, or changing treatments.

Diffusion basis spectrum imaging (DBSI) utilizes a data-driven multiple-tensor modeling approach to differentiate coexisting morphological features resulting from tumor pathologies or other attributes within an image voxel. We have previously demonstrated that DBSI quantifies tissue injury in an array of central nervous system disorders including multiple sclerosis (13-15), cervical spondylotic myelopathy (16), and epilepsy (17). In this study, we demonstrate both Gd-enhanced T1WI and hyperintense FLAIR areas contain a spectrum of DBSI-derived morphological signatures in GBM. Using a modified DBSI algorithm to separate inflammation from tumor cellularity, we show DBSI-derived restricted-isotropic-diffusion fraction positively correlated with tumor cellularity in GBM specimens. Finally, to improve the performance of a DBSI-based cancer detection, we developed a robust Diffusion Histology Imaging (DHI) approach, combining a machine learning algorithm and DBSI metrics, to accurately identify and classify various histopathological components of GBM.

## Materials and Methods

### Study Design

This study was approved by the Institutional Review Board of the Washington University School of Medicine (St. Louis, Missouri) and conducted in accordance with the Declaration of Helsinki. Written informed consents were obtained from all participants. The goal was to develop a reliable and consistent neuroimaging outcome measure for accurately classifying high cellularity tumor, tumor necrosis and tumor infiltration in high grade glioma. The inclusion criteria: (i) adult subjects scheduled for brain tumor resection at the Washington University School of Medicine, (ii) subjects had not received radiation therapy or chemo therapy, and (iii) subjects whose resected tumor specimen sufficiently large for *ex vivo* MRI in addition to that required for clinical diagnosis. Sixteen newly-diagnosed adult GBM patients (Table S1) were recruited for *in vivo* (n = 3) and *ex vivo* (n = 13) MRI studies from June 2015 to January 2017. Eighteen patients with GBM suspicion were recruited for eligibility assessment from June 2015 to January 2017 (Fig. S1). Two patients were excluded due to data problem and pathalolgocial assessment of non-GBM (anaplastic ependymoma), respectively. After exclusion, sixteen newly-diagnosed adult GBM patients without any previous treatments were included for *in vivo* (n = 3) and *ex vivo* (n = 13) MRI studies. The patient characteristics were summarized in Table S1.

### Surgical Resection of Brain Tumor Specimen

Nineteen resected specimens from 13 GBM patients underwent multi-slice/section *ex vivo* MRI and histological examinations (Fig. S1). The average size of the specimens was 8 mm ± 4 mm. Among the 19 specimens, at least one specimen was obtained from each individual; for three individuals, specimens were taken from two sectors in the tumor; and from one individual, specimens were taken from four tumor sectors. Each tissue specimen contains multiple image slices and histology sections for MRI-histology co-registration and quantification. Multi-slice/section DBSI and H&E revealed that patterns of GBM pathologies were similar throughout the thickness of all specimens with the exception of two specimens from one individual, in which H&E patterns were distinct in two sections. Thus, we analyzed a total of 21 DBSI-H&E matched sections from 19 specimens.

### Sample preparations

After resection, specimens were immediately fixed in 10% formalin in phosphate buffered saline (PBS, PH = 7.4) at room temperature for at least 48 hours (Fig. S2A) and then transferred to PBS. PBS was changed every two days for a total of two weeks before the experiment. The specimens were embedded in agar gel for MR imaging and then analyzed using DBSI and DTI.

### *Ex Vivo* MRI of Surgical Resection Tumor Specimens

Specimens were formalin-fixed at time of collection and then agarose-gel-embedded (Fig. S2B) before being examined using a 4.7-T Agilent MR scanner (Agilent Technologies, Santa Clara, CA) with a home-made circular surface coil (1.5-cm diameter, Fig. S2C). A multi-echo spin-echo diffusion-weighting sequence with 99 diffusion-encoding directions (maximum b-value at 3000 s/mm^2^) was employed to acquire DWI with a 0.25 × 0.25 mm^2^ in-plane resolution, and 0.5-mm thickness. The imaging parameters were as follows: repetition time (TR) 1500 ms, echo time (TE) 40 ms, time between application of gradient pulse 20 ms, diffusion gradient on time 8 ms, slice thickness 0.5 mm, field-of-view 24 × 24 mm^2^, data matrix 96 × 96, number of average 1, in-plane resolution 0.25 × 0.25 mm^2^. Total acquisition time was approximately 4 hours. MR images were zero-filled to 0.125 × 0.125 mm^2^ in-plane resolution for DBSI and diffusion tensor imaging (DTI) analyses.

### Histological Sectioning and Staining

The specimens underwent sequential sectioning at 5-μm thickness. Sections were individually stained with hematoxylin and eosin (H&E) and glial fibrillary acidic protein (GFAP). Histology slides were digitized using NanoZoomer 2.0-HT System (Hamamatsu, Japan) with a 20× objectives for analyses. Each tissue specimen contains multiple image slices and histology sections for MRI-histology co-registration and quantification.

### Cellularity Quantified in H&E and GFAP

We developed a procedure involving down-sampling histological images and co-registering MRI with histological images to enable the voxel-wise correlation between tumor cellularity and MRI-derived surrogate marker of cellularity, e.g., DTI-derived ADC and DBSI-derived restricted diffusion fraction. Detailed analysis was performed as described in Supplementary Materials and Methods. Briefly, specimens were sectioned and stained after *ex vivo* MRI to acquire H&E and GFAP images. High resolution histology images were down-sampled to match DBSI/DTI resolution (125 × 125 µm^2^) to enable direct comparison between DBSI/DTI and histological images. Each down-sampled histology image voxel contained 272 × 272 original image pixels. A two-dimensional (2D) thin-plate-spline (TPS) co-registration method was adopted using 30 manually-picked landmarks.

### *In Vivo* MRI of Human Subjects

A 3-T Siemens TIM Trio (Erlanger, Germany) with a 32-channel head coil was used for all *in vivo* images. Axial diffusion-weighted images (DWI) covering the whole brain were acquired using a multi-b value diffusion weighting scheme (99 directions, maximum b-value 1500 s/mm^2^) and the following parameters: TR = 10,000 ms; TE = 120 ms; FOV = 256 × 256 mm^2^; slice thickness = 2 mm; in-plane resolution = 2 × 2 mm^2^; total acquisition time = 15 min. Eddy current and motion artifacts of DWI were corrected before susceptibility-induced off-resonance field was estimated and corrected. Conventional MRI sequences including gadolinium-enhanced T1-weighted image, anatomical 3D magnetization prepared rapid acquisition of gradient echo (MPRAGE) image, T2-weighted image and T2-weighted fluid attenuated inversion recovery (FLAIR) image were performed per standard clinical protocol.

### Recapitulating Neuropathological Analysis of GBM

Pathological examination following stereotactic biopsy or surgical resection plays a vital role in current clinical decision-making for the management of GBM patients, based on the neuropathologist’s recognition of morphological signatures reflecting tumor cells and changes in the microenvironment, including treatment effects, which are characteristics missed by current MRI biomarkers. To address this critically important unmet need, we developed diffusion basis spectrum imaging (DBSI), which utilizes a data-driven multiple-tensor modeling approach to disentangle pathology and structural profiles within an image voxel (13,18-22). Although DBSI-derived structural metrics distinguish and quantify various tissue pathologies in an array of CNS disorders (13,19,23-26), the ability of DBSI to detect tissue microstructure alone is insufficient to accurately identify the underlying GBM pathologies of high tumor cellularity, tumor necrosis, and tumor-infiltrated white matter. Thus, we have developed a novel *Diffusion Histology Imaging (DHI) approach*, which applies machine/deep learning algorithms (27,28) using DBSI structural metrics as the input classifiers, to accurately model underlying GBM pathologies.

DBSI models brain tumor diffusion-weighted MRI signals as a linear combination of discrete multiple anisotropic diffusion tensors and a spectrum of isotropic diffusion tensors:

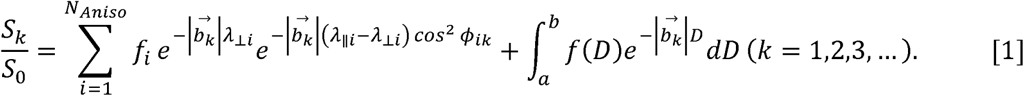

Here *b*_*k*_ is the *k*^th^ diffusion gradient. *S*_*k*_*/S*_*0*_ is the acquired diffusion-weighted signal at direction of *b*_*k*_ normalized to non-diffusion-weighted signal. *N*_*Aniso*_ is number of anisotropic tensors to be determined. *ϕ*_*ik*_ is the angle between diffusion gradient *b*_*k*_ and principal direction of the *i*^*th*^ anisotropic tensor. 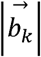 is *b*-value of the *k*^*th*^ diffusion gradient. *λ*_*||i*_ and *λ*_*⊥i*_ are axial and radial diffusivity of the *i*^*th*^ anisotropic tensor under the assumption of cylindrical symmetry; *f*_*i*_ is signal-intensity-fraction of the *i*^*th*^ anisotropic tensor. *a* and *b* are low and high diffusivity limits of isotropic diffusion spectrum. *f(D)* is signal-intensity-fraction at isotropic diffusivity *D*.

Based on our *ex vivo* MRI and histological analyses of resected specimens, the following isotropic-diffusion profiles have been established based on diffusivity. We observed that highly restricted isotropic diffusion (0 ≤ D ≤ 0.2 μm^2^/ms) is associated with lymphocytes; restricted-isotropic diffusion (0.2 < D ≤ 0.8 μm^2^/ms) is associated with high tumor cellularity in GBM; and hindered-isotropic diffusion (0.8 < D ≤ 2 µm^2^/ms) is associated with tumor necrosis. For *in vivo* human subjects, the *in vivo* diffusivity profile can be estimated by extrapolating *ex vivo* diffusivity based on the temperature difference: highly restricted diffusion fraction (0 ≤ D ≤ 0.2 μm^2^/ms; not affected by temperature); restricted isotropic diffusion (0.2 < D ≤ 1.5 µm^2^/ms); and hindered isotropic diffusion (1.5 < D ≤ 2.5 µm^2^/ms). Further detailed information can be found in the supplementary methods.

DBSI provides a simple tensor expression to visualize morphological features resulting from tumor formation and non-tumor entities appearing indistinct to tumor by conventional MRI. For example, in an image voxel where normal white matter tracts and gray matter is coexisting with the presence of tumor cells. The tensor function of anisotropic and isotropic tensors will not change comparing with the normal tissues with the exception of changes resulting from the presence of tumor cells. If necrosis is present in tumor containing image voxels, it would require multiple isotropic diffusion tensors, such as restricted (modeling tumor cells) and hindered (modeling necrosis) isotropic tensors, to completely model the diffusion-weighted signals. The different tensor expressions of individual image voxels thus bear morphological signatures of underlying pathology. In the case where tumor cells happen to also damage white matter tracts resulting, say, axonal injury and demyelination. The isotropic tensors within this image voxel will remain the same but now anisotropic diffusion tensor will exhibit decreased axial diffusivity and increased radial diffusivity. It is the sensitivity of diffusion-weighted MRI signal to the microstructural changes in the scale up to 10 µm range (depending how one adjusts diffusion-weighting condition) that allows DBSI to more precisely reflect morphological changes resulting from tumor presence or other pathological conditions. By taking the advantage of this feature of DBSI as the input of machine learning algorithms, we created DHI to recapitulate histopathological analysis using MRI.

### Statistical Analysis

We used Spearman’s rank correlation to measure strengths of monotonically increasing or decreasing associations between histology and MRI cellularity measurements. Statistically significant results were determined at a pre-determined alpha level of 0.05.

To construct a machine learning classifier for histopathological prediction, support vector machine (SVM) with polynomial kernel algorithm was adopted using a package by Scikit-learn in Python (29). We performed a linear co-registration between MRI and corresponding neuropathologist-classified H&E images to label each image voxel with the gold standard of pathology. Total 21 sections from 19 brain tumor specimens (6,605 image voxels) were analyzed to determine DBSI and DTI metric profiles of each image voxel. Image voxels from four randomly selected sections were used for testing and the voxels from remaining sections were used for training. For cross validation purpose, total 1000 distinct training-test group pairings were run to prevent selection bias. Additionally, to address potential internal correlations of voxels from same patients, we performed 500 random splits that assigned voxels from different patients into training and test datasets. Mean values and 95% confidence intervals were calculated.

Confusion matrices were calculated to illustrate the specific examples of tumor pathologies where predictions were discordant with pathologist-identified pathologies. We evaluated overall classification accuracy of testing voxels as well as true positive rate, true negative rate and positive predictive value of the model prediction. Receiver operating characteristics (ROC) and precision-recall curves were calculated using a one-versus-rest strategy to assess model discrimination for each tumor pathology. Area under curves (AUC) and F_1_-scores were calculated to compare the relative performance of DHI to pathologist-identified pathologies.

## Results

### Patient Information

Among the sixteen patients, eleven were male and five were female. The mean age at diagnosis was 61.1 ± 14.2 years. Pathological analysis of tumor specimens confirmed isocitrate dehydrogenase*-*wildtype GBM in all 16 patients (Table S1).

### DBSI Metrics Are Not Unique to Gd-Enhanced, Non-Enhanced T1W or Hyperintense FLAIR Tumor Regions in Patients

We performed clinical MRI and DBSI on three GBM patients. Representative Gd-enhanced T1W, FLAIR, T2W, DBSI, and ADC images were obtained from a 79-year-old male patient with a right temporal GBM (Fig. 1A). We outlined Gd-enhanced and non-enhanced T1W regions to compare the underlying DBSI metrics in these regions, overlaid on MPRAGE-T1W images (Fig. 1A). Based on our previous DBSI applications, we predict that DBSI metrics of restricted fraction, hindered fraction, and anisotropic fraction, would be seen in regions/voxels containing high tumor cellularity, necrosis, and fiber-like structures (neuronal fibers or extracellular matrix fibers), respectively. Strikingly, DBSI metrics of restricted fraction (red), hindered fraction (blue), and anisotropic fraction (green) were entangled in both Gd-enhancing and non-enhancing regions (Fig. 1A, DBSI). Specifically, hyperintensity of restricted fraction were widespread in Gd non-enhancing region where histology is typically considered to be necrosis, indicating the potential high tumor cellularity in this region that could significantly challenge the current clinical standard.

**Fig. 1.**
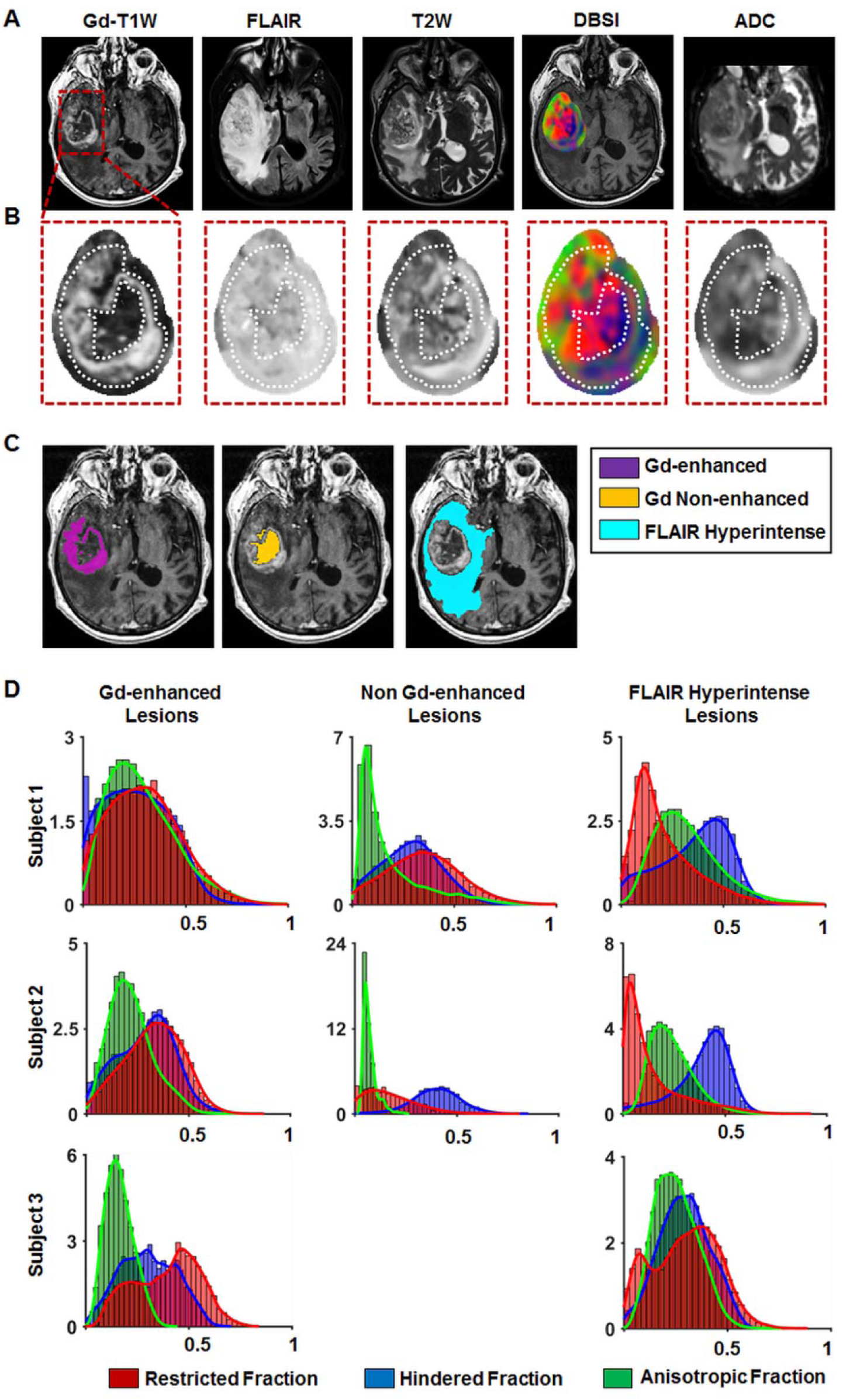
Gd-enhanced T1WI or hyper-intense FLAIR failed to reflect the morphological complexity underlying GBM. One rim enhancing lesion in Gd-T1WI at the right temporal lobe of a 79-year-old male patient was identified (A, red square) and enlarged (B, red square). The exact same regions from FLAIR, T2WI, DBSI and ADC were also displayed for comparisons (B, red squares). Gd enhanced region within this lesion was further outlined in the Gd-T1WI and applied to other images (B, white dash outlines). Both Gd-enhanced and non Gd-enhanced regions exhibit various extents of restricted diffusion (red), hindered diffusion (blue), and anisotropic diffusion (green), suggesting the lack of pathological or structural specificity of the widely used Gd-enhanced T1W and hyper-intense FLAIR lesion. Contrary to widely accepted notion that Gd-enhanced T1WI lesion is primarily associated with tumor cellularity, we observed the elevated putative DBSI cellularity marker (restricted fraction; red, scale 0 - 0.6) in both Gd-enhanced and non Gd-enhanced regions. Putative tumor necrosis or tissue loss (hindered fraction; blue, scale 0 - 1.0) is also seen in both regions. DBSI anisotropic diffusion fraction (reflecting the fiber volume fraction of neuronal fibers or collagen fibers; green, scale 0 - 1.0) is also present in both Gd-enhanced and non Gd-enhanced regions (within rim enhanced area). Hyper-intense Gd-T1W lesions (C, purple mask), hypo-intense Gd-T1W lesions (C, yellow mask), and hyper-intense FLAIR lesions (C, cyan mask) were segmented to quantitatively analyze the histogram of DBSI metrics one these three types of lesions from three GBM patients (D; x-axis = fraction of DBSI metric; y-axis = number of occurrence). As seen in these three subjects (D, subject 3 does not have a non Gd-enhanced lesion), the three DBSI metrics are present in all Gd-enhanced, non Gd-enhanced and FLAIR hyperintense lesions, further supporting the insufficiency of these commonly employed imaging markers.

To determine if specific DBSI structural metrics are enriched in particular clinical MRI sequences, we generated histograms of DBSI-metrics from Gd-enhancing, non-enhancing, and FLAIR hyperintense lesions from all three patients (Fig. 1B). The common feature among the three GBM cases was the consistent presence of the three DBSI metrics in all clinical MRI-defined lesions. Qualitatively, the pattern of DBSI metric distributions did not appear to be unique for Gd-enhancing, non-enhancing, or FLAIR hyperintense regions of GBM tumors, suggesting that these clinical MRI-defined regions harbor mixed pathologies.

### Tumor Cellularity Correlated with DBSI-Restricted Fraction, but Not ADC, in *Ex Vivo* GBM Specimens

As shown above (Fig. 1), *in vivo* DBSI restricted, hindered, and anisotropic fractions were highly overlapping in MR-lesions of GBM. To definitively determine relationships between DBSI metrics and GBM pathologies, we examined *ex vivo* DBSI metrics in histologically-identified regions of high tumor cellularity, tumor necrosis, and tumor infiltration in 19 surgically-resected specimens. We performed a thin-plate-spline co-registration on specimens correlating diffusion-weighted images with H&E and glial fibrillary acidic protein (GFAP) cellularity maps (Fig. 2A/B) to allow voxel-to-voxel correlation of histology (H&E and GFAP positive area ratio maps) with ADC, and DBSI-restricted fraction maps (Fig. 2C). We randomly selected fifty voxels from down-sampled H&E images (Fig. 2A, red squares) and mapped them to the co-registered GFAP, MRI-metric maps for voxel-based correlation. Out of twenty specimens, fifteen underwent MRI-H&E and nine underwent MRI-GFAP correlation analyses. The rest were excluded due to unmatched sectioning planes.

**Fig. 2.**
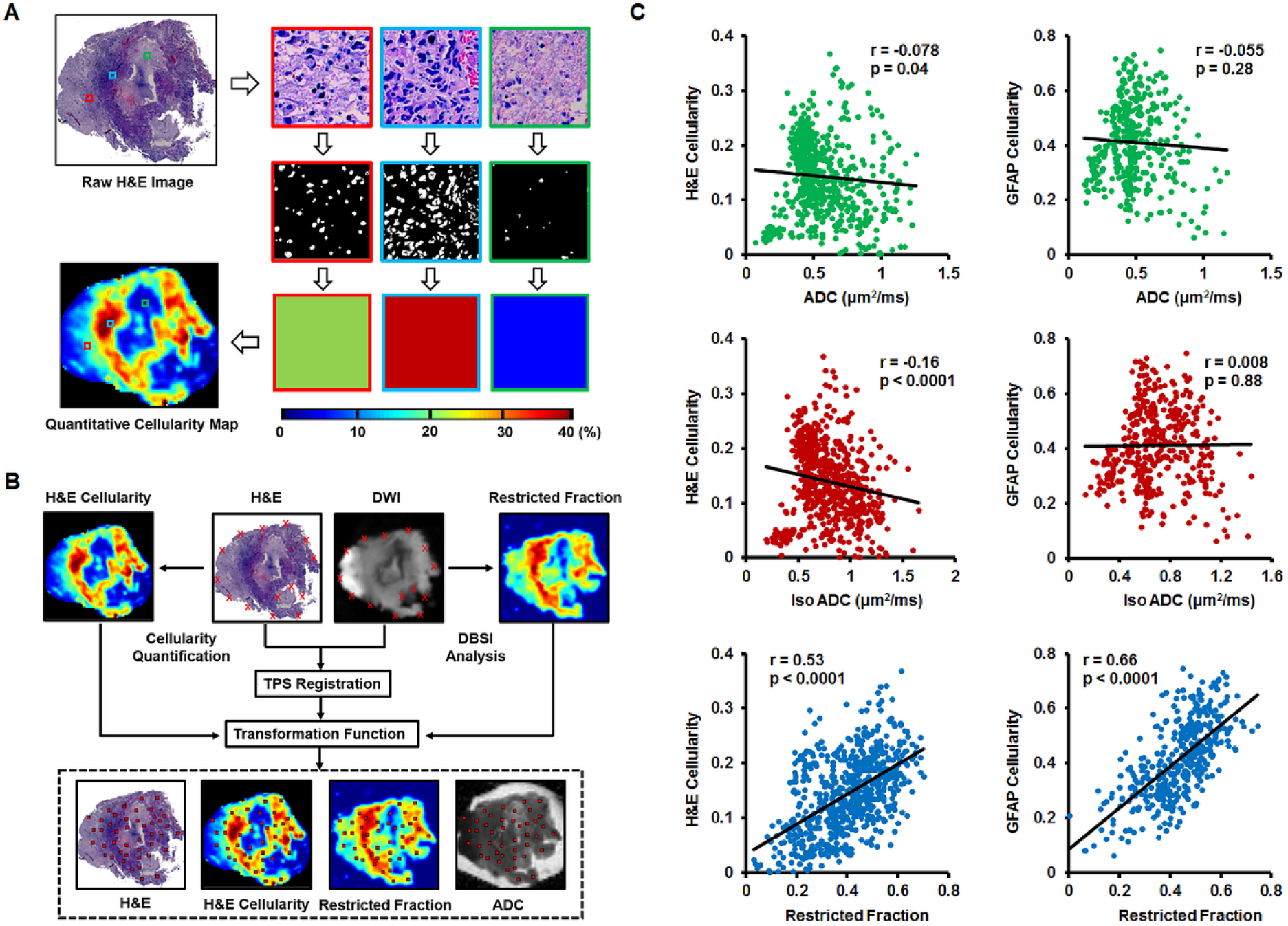
MRI-Histology co-registration and quantification. **(A)** Quantitative cellularity maps were calculated from high resolution H&E images. High-resolution H&E images were down-sampled to match MRI resolution (125 × 125 μm^2^). Individual tiles of MRI voxels containing 272 × 272 high-resolution H&E image pixels were extracted. Fractions of positively-stained area of individual image tiles were computed from ratios between positively-stained areas and total pixel areas and color-coded then stitched back to construct the quantitative cellularity map. **(B)** We performed co-registration of DWI and H&E images to allow voxel-to-voxel correlation of histology with ADC, DBSI-isotropic ADC, and DBSI-restricted fraction. Around thirty landmarks were manually placed along the perimeter of diffusion-weighted images and down-sampled histology images for co-registration. The transformation function of thin-plate-spline co-registration was applied to warp MR images to match histology images. Fifty image voxels were randomly selected from each down-sampled H&E image and applied to all co-registered maps for correlation and quantitative analysis. **(C)** Regression analysis of DTI-ADC vs. H&E and DTI-ADC vs. GFAP suggested weak correlations (r = -0.078, -0.055 and p = 0.04, 0.28, respectively). DBSI isotropic-ADC showed the expected negative correlation with H&E-cellularity (r = -0.16, p < 0.0001) but did not correlate with GFAP-cellularity (r = 0.008, p = 0.88). DBSI-restricted fraction displayed statistically significantly high correlation with H&E- and GFAP-cellularity (r = 0.53, 0.66, respectively; p < 0.0001).

Spearman’s rank correlation for selected voxels from all specimens (Fig. 2C) was used to assess the general performance of imaging biomarkers for cellularity in tumor samples. Restricted fraction correlated with H&E (r = 0.53, p < 0.0001) and GFAP positive areas (r = 0.66, p < 0.0001). In contrast, ADC failed to correlate with H&E (r = -0.078, p = 0.04) or GFAP (r = -0.055, p = 0.28). Additionally, DBSI isotropic-ADC showed slightly negative correlation with H&E-cellularity (r = -0.16, p < 0.0001) and no correlation with GFAP-cellularity (r = 0.008, p = 0.88).

### Qualitative Comparison of DBSI Metrics with GBM Pathologies

To definitively determine relationships between DBSI metrics and GBM pathologies, we examined *ex vivo* DBSI metrics in histologically-identified regions of high tumor cellularity, tumor necrosis, and tumor infiltration in 19 surgically-resected specimens. A representative tumor specimen (10.1 × 8.7 mm^2^) from a 77-year-old female patient demonstrates the relationship between DBSI metrics and tumor pathologies (Fig. 3). Hyperintense DWI (*i.e.* hypointense ADC) defined voxels did not correspond to H&E measures of cellularity (Fig. 2 and 3A/B). By co-registering DWI with histological images, representative voxels from regions of high tumor cellularity (Fig. 3B, H&E & GFAP; red square), tumor infiltration (Fig. 3B, H&E & GFAP; green square), and tumor necrosis (Fig. 3B, H&E & GFAP; blue square) were selected to perform isotropic diffusion spectrum analysis (Fig. 3C). These representative image voxels of H&E and GFAP were enlarged and displayed for validating the histological findings (Fig. 3B). In general, the isotropic diffusion spectrum analysis indicated three distinct clusters of diffusion signatures. Specifically, high tumor cellularity areas exhibited peaks at highly-restricted and restricted diffusion regions of DBSI-isotropic diffusion spectrum; infiltrated white matter exhibited peaks at the same locations as high cellularity with varying intensities; and tumor necrosis exhibited highly-restricted and hindered diffusion peaks (Fig. 3C). Based on the diffusion spectrum analysis from each image voxel, the diffusion tensor fraction maps could be derived. Although these DBSI metrics were overlapping in these tumor pathologies (Fig. 1A/B and 3C), DBSI restricted and hindered fraction maps qualitatively resembled areas of high tumor cellularity and tumor necrosis, respectively, as identified by a neuropathologist (Fig. 3D).

**Fig. 3.**
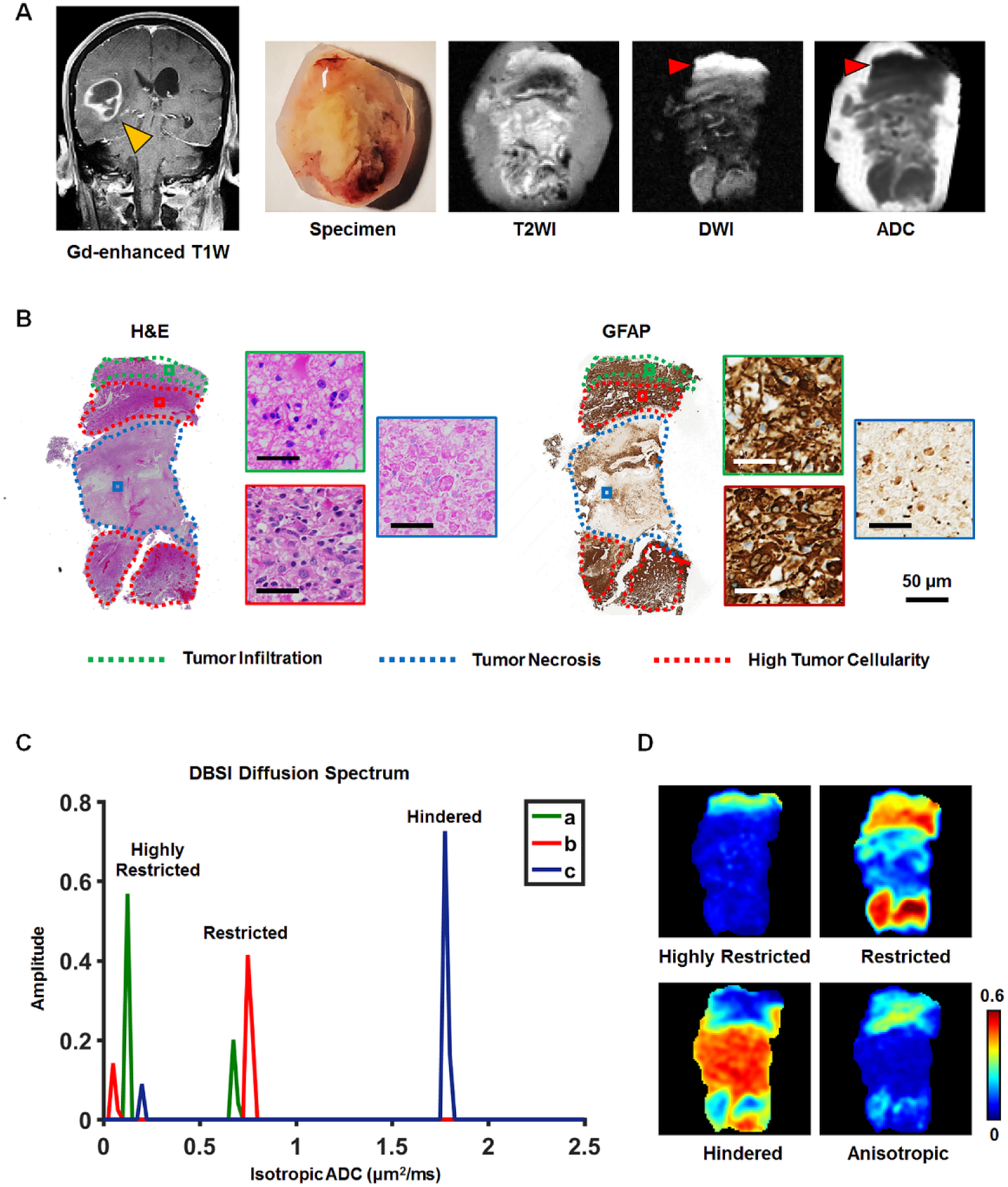
Association between DBSI-metrics and neuropathologist-identified tumor pathology. A surgically-resected specimen from a 77-year-old female GBM patient was analyzed via T2WI and DWI (A). Neuropathologist identified high tumor cellularity, tumor infiltration, and tumor necrosis regions in H&E and GFAP staining slides and digitized images (B). According to the widely accepted notion, a hyper-intense DWI (red arrow), i.e., hypo-intense ADC (red arrow), region is suggestive of increased tumor cellularity. However, it does not match neuropathologist-identified pathology featuring white matter tracts with tumor infiltration based on histology staining (B), consistent with figure 2 findings. From the co-registered MRI-histology images, high tumor cellularity, tumor infiltration, and tumor necrosis regions were matched with DBSI-metrics. High tumor cellularity signal (red) exhibits peaks at highly-restricted and restricted diffusion regions; infiltrated white matter signal (green) exhibits peaks at the same locations as high cellularity with varying intensities; and tumor necrosis signal (blue) exhibits highly-restricted and hindered diffusion regions (C). Based on these distributions, we generated DBSI highly-restricted, restricted, and hindered isotropic-diffusion signal fraction maps (D). These maps reveal that highly-restricted fraction is high in tumor infiltration and high tumor cellularity regions; restricted fraction is highly associated with high tumor cellularity regions (consistent with findings of figure 2); and hindered diffusion fraction is highly correlated with H&E tumor necrosis regions. The intensity gradient on restricted fraction map reflects tumor cellularity change.

### Accurate Prediction of Pathological Features in GBM Using Diffusion Histology Imaging

Through image co-registration, MRI voxels corresponding to pathologically-verified areas of high tumor cellularity, tumor necrosis, and tumor infiltration were identified. Image voxel values of DTI (Fig. 4A) and DBSI (Fig. 4B) metrics are presented to demonstrate the distinctions and similarities among these identified tumor pathologies. A multi-parametric examination based on restricted fraction, hindered fraction, and isotropic fraction separated the three pathologically distinct entities (Fig. 4C), suggesting analysis based on multiple DBSI metrics could potentially better distinguish among these pathologies rather than single DBSI metrics alone.

**Fig. 4.**
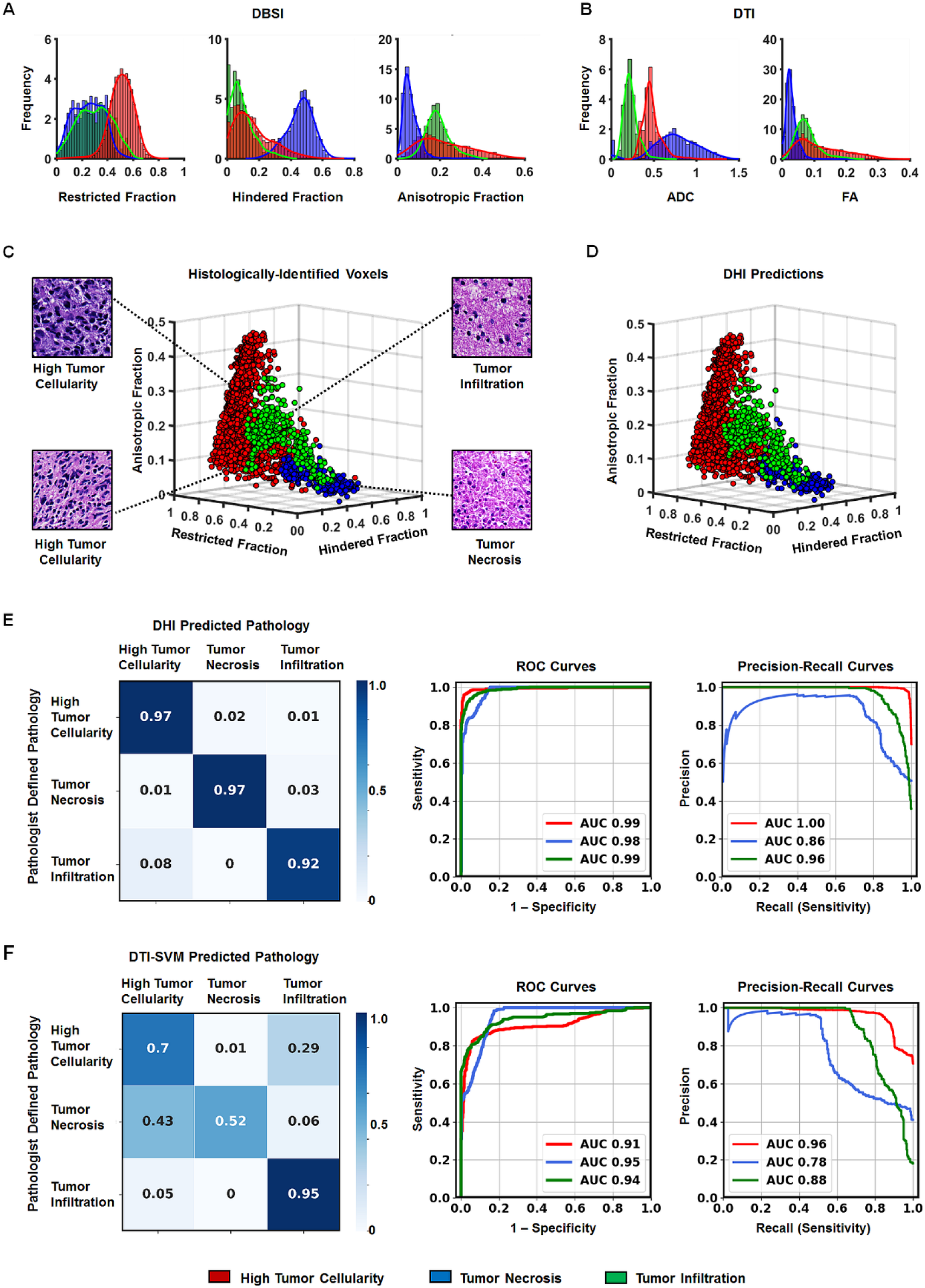
Classifying high tumor cellularity, tumor necrosis and tumor infiltration in resected GBM specimens. The structural metrics derived from DBSI (A) and DTI (B) were obtained in neuropathologist-identified high tumor cellularity (red), tumor necrosis (blue), and tumor infiltration (green) regions through MRI-histology co-registration. Overlapping profiles of DTI/DBSI structural metrics are common within individual tumor pathology. Thus, it is difficult to distinguish tumor pathologies by thresholding single diffusion metrics values. Representative neuropathologist-identified histology-image voxel values of DBSI restricted, hindered, and anisotropic fractions reveal that the three tumor pathologies can be resolved by combining the three DBSI metrics (C). Representative histology images corresponding to selected DBSI image voxels were presented. For this independent dataset (n = 1,963), DHI predicted voxels showed great match with histology, affording a 96.2% overall accuracy in predicting high tumor cellularity, tumor necrosis and tumor infiltration (D). Confusion matrices reveal DHI (E) is more accurate than DTI-SVM (F) in predicting high tumor cellularity, tumor necrosis and tumor infiltration. Additionally, DHI showed greater AUC values than DTI-SVM on both ROC and precision-recall curves.

We thus developed DHI incorporating a supervised SVM algorithm with modified-DBSI (incorporating a distinction between inflammation and tumors) derived structural metrics as classifiers to construct predictive models to distinguish among different tumor histopathology. We trained and validated the predictive model on image voxels from 17 of the 21 GBM specimen sections. The established model was applied to image voxels from four remaining GBM specimen sections to predict distributions of high tumor cellularity (Fig. 4D, red), tumor necrosis (Fig. 4D, blue) and tumor infiltration (Fig. 4D, green) with 96.2% overall accuracy (n = 1963). DHI correctly predicted 97.2%, 96.6% and 91.8% of the image voxels as high tumor cellularity, tumor necrosis, and tumor infiltration, respectively.

A comparison between DHI and DTI-SVM was performed using confusion matrices. The DHI (Fig. 4E) approach demonstrated better prediction accuracy for tumor pathologies compared to DTI-SVM (Fig. 4F). We also performed ROC and precision-recall curves analyses for each tumor pathological feature (Fig. 4E, F). DHI indicated greater ROC and precision-recall AUC values for all the pathological features than DTI-SVM.

### Pathological Validation of DHI

From four DHI test specimens, we randomly selected four DWI voxels from each specimen (Fig. 5A - D) and use corresponding histology as validation. This was achieved by tracking each voxel back to the co-registered down-sampled histological images to compare DHI-predicted pathologies with gold standards. We observed high predictive performance of DHI on individual specimens.

**Fig. 5.**
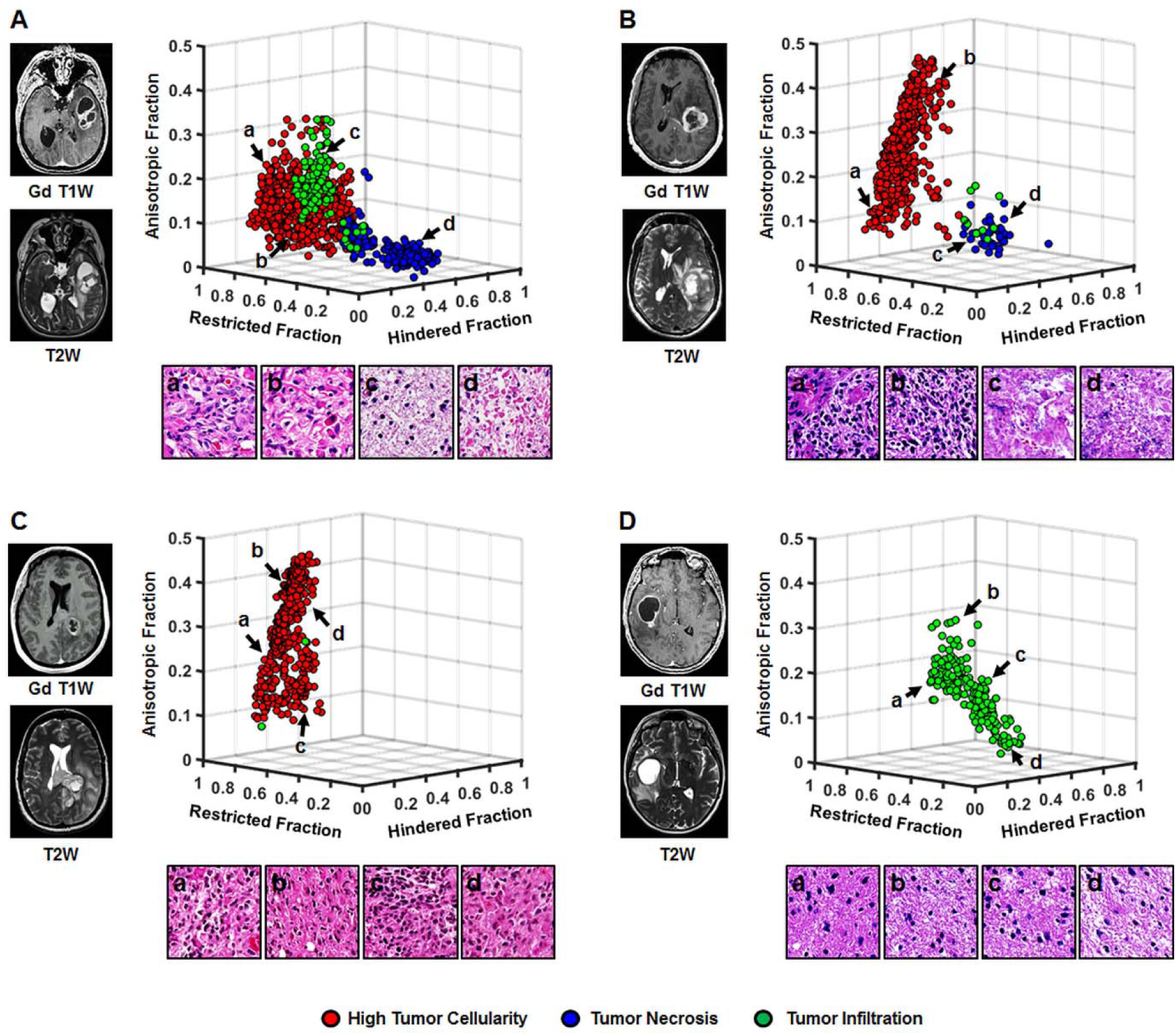
Histology validation of DHI determined tumor pathologies in the four test specimens. In a 77-year-old female GBM patient (B122) specimen, DHI correctly predicts high tumor cellularity (A, red), tumor necrosis (A, blue) and tumor infiltration (A, green) with 94.3%, 97.3% and 82.1%, respectively. Corresponding H&E image tiles verify the randomly-selected DHI-determined high tumor cellularity (A: a, b), infiltration (A: c) and necrosis (A: d). The second test specimen from a 54-year-old male GBM patient (B95) exhibits a 98.0% and 93.3% true prediction rate of DHI-determined high tumor cellularity and necrosis voxels, respectively, validated by corresponding H&E image tiles: high tumor cellularity (B: a, b) and tumor necrosis (B: c, d). The third specimen from a 47-year-old female GBM patient (B128) was also assessed to reveal that DHI-determined high tumor cellularity is 99.0% accurate, as validated by the H&E tiles (C: a, b, c, d). In the fourth test specimen from a 57-year-old GBM female patient (B94), DHI correctly predicted 100% of the tumor infiltration voxels (D). All the four selected voxels from co-registered H&E (D: a, b, c, d) indicated tumor infiltration pattern.

DHI predicted 94.3% of high tumor cellularity areas (Fig. 5A, red), 97.3% of necrotic areas (Fig. 5A, blue) and 82.1% of tumor infiltration areas (Fig. 5A, green) in a 77-year-old female patient specimen (B122). Indeed, corresponding H&E image tiles (i.e., voxels of down-sampled histology images) verified the randomly-selected DHI predictions of high tumor cellularity (Fig. 5A: a, b), tumor infiltration (Fig. 5A: c), and tumor necrosis (Fig. 5A: d). The second test specimen from a 54-year-old male patient (B95) exhibited 98.0% accuracy of DHI predictions of high tumor cellularity voxels (Fig. 5B, red) and 93.3% true prediction rates of tumor necrosis (Fig. 5B, blue) as validated by corresponding H&E image tiles (high tumor cellularity (Fig 5B: a, b) and tumor necrosis (Fig 5B: c, d). The third specimen from a 47-year-old female patient (B128) was also assessed, demonstrating that DHI predicted voxels of high tumor cellularity was 99.0% accurate (Fig. 5C: a, b, c, d). In the fourth test specimen from a 57-year-old female patient (B94), DHI correctly predicted 100% of tumor infiltration voxels (Fig. 5D). Corresponding H&E image tiles all indicated tumor infiltration patterns.

### Comparing DTI-SVM and DHI Performance on Predicting Tumor Pathologies

We ran 1000 random training/validation and test split pairings to address possible selection bias resulting from the use of specific test samples. The mean accuracy of DHI was 89.6%, compared to 76.7% of DTI-SVM. Mean true prediction rates of DHI for high tumor cellularity, tumor necrosis, and tumor infiltration were 87.5%, 89.0% and 93.4%, respectively (Table 1). By contrast, mean true prediction rates of DTI-SVM were 76.7%, 62.3% and 97.9%, respectively (Table S2). Additionally, DHI showed much better overall precision-recall performances, with mean F_1_-scores of DHI as 0.917, 0.823 and 0.876 for three tumor pathologies. We further performed ROC analyses using one-vs.-rest strategy to test how well our classifiers to distinguish one tumor pathology from others (e.g., infiltration vs. non-infiltration). The ROC analysis results revealed that DHI had great performance on distinguishing these three tumor pathologies, with mean AUC values of 0.975, 0.989 and 0.951 for high tumor cellularity, tumor necrosis, and tumor infiltration, respectively.

**Table 1.** Diagnostic Performances of DHI on Predicting Tumor Pathologies. Values were summarized as mean (95% CI). Sample-wise split models assigned voxels from different samples into training-validation dataset and test dataset. Patient-wise split assigned voxels from different patients into train-validation dataset and test dataset. AUC, area under curve. CI, confidence interval.

|  | **Tumor Pathology** | **Sensitivity (%)**<br>**(95% CI)** | **Specificity (%)**<br>**(95% CI)** | **AUC**<br>**(95 CI)** | **F<sub>1</sub>-Score**<br>**(95% CI)** |
| --- | --- | --- | --- | --- | --- |
| Sample-wise split | High Tumor Cellularity | 87.5<br>(86.7 □ 88.3) | 95.3<br>(94.9 □ 95.8) | 0.975<br>(0.973 □ 0.976) | 0.917<br>(0.911 □ 0.923) |
|  | Tumor Necrosis | 89.0<br>(88.3 □ 89.8) | 92.9<br>(92.5 □ 93.4) | 0.951<br>(0.947 □ 0.955) | 0.823<br>(0.814 □ 0.833) |
|  | Tumor Infiltration | 93.4<br>(92.7 □ 94.0) | 95.9<br>(95.5 □ 96.3) | 0.989<br>(0.987 □ 0.990) | 0.876<br>(0.865 □ 0.887) |
| Patient-wise split | High Tumor Cellularity | 85.5<br>(84.5 □ 86.4) | 92.0<br>(91.5 □ 92.5) | 0.960<br>(0.958 □ 0.963) | 0.893<br>(0.887 □ 0.900) |
|  | Tumor Necrosis | 93.7<br>(93.1 □ 94.3) | 90.6<br>(90.1 □ 91.2) | 0.938<br>(0.933 □ 0.942) | 0.816<br>(0.810 □ 0.822) |
|  | Tumor Infiltration | 81.4<br>(79.9 □ 82.8) | 96.9<br>(96.4 □ 97.3) | 0.989<br>(0.988 □ 0.990) | 0.816<br>(0.806 □ 0.825) |

To address the potential internal correlations from voxels of the same patients in training/validation and test datasets, we performed 500 random training/validation and test splits that assigned voxels from different patients into training/validation and test datasets. As expected, the accuracy was slightly lower, with 87.1% compared to 89.6% from the sample-wise split method. Similarly, the AUCs (high tumor cellularity: 0.960; tumor necrosis: 0.938; tumor infiltration: 0.989) and F_1_-scores (high tumor cellularity: 0.893; tumor necrosis: 0.816; tumor infiltration: 0.816) were also slightly lower but comparable to results from sample-wise split method (Table 1), indicating the consistency and generalization of DHI.

## Discussion

The standard of care for GBM involves surgical resection, followed by radiotherapy with concurrent and adjuvant chemotherapy. Histological assessment of tumor cellularity, necrosis, and infiltration plays a vital role in the clinical decision-making for the management of GBM patients. The current gold standard of pathological examination following stereotactic biopsy or surgical resection (30) carries potential risks (31). On some occasions, inconclusive pathological findings may result from inadequate sampling that may necessitate repeat procedures (32). Thus, noninvasive neuroimaging approaches to facilitate diagnosis or to guide biopsies and surgical planning are needed to improve GBM patient care.

Through voxel-wise comparisons with histological images, we demonstrated that DBSI-derived restricted-isotropic-diffusion fraction, hindered-isotropic-diffusion fraction, and anisotropic-diffusion-fraction closely correlate with high tumor cellularity, tumor necrosis, and fiber-like structures, respectively. However, these metrics alone were not sufficient to clearly distinguish high tumor cellularity, tumor necrosis, or tumor infiltration (Figs. 1, 3 and 4). We thus developed DHI, incorporating an SVM predictive model using DBSI metrics as the classifiers, to successfully predict high tumor cellularity, tumor necrosis, and tumor infiltration against the gold standard of histology with high accuracy (Figs. 4 - 5).

Various neuroimaging techniques have been tested to assess the treatment response of brain tumors in clinical practice. Among the wide range of available neuroimaging modalities, contrast-enhanced T1W image is currently the method of choice for brain tumor diagnosis. Unfortunately, Gd-enhanced T1W image lacks specificity because it merely reflects a disrupted blood-brain barrier (33). Chemotherapy, radiation, and newer clinical trial treatments such as immunotherapies produce neuroimaging lesions that mimic tumor progression or recurrence, further confounding clinical decision making (34). These and other shortcomings of current clinical MRI sequences suggest limitations of the MacDonald criteria (4) and the Response Assessment in Neuro-Oncology (RANO) updated response assessment criteria (3,35) in monitoring tumor burden. Therefore, there is an urgent need to develop imaging modalities that can non-invasively detect and characterize the histological features of post-treatment GBM for appropriate treatment planning.

Advanced MRI methods, such as perfusion-weighted imaging with and without contrast (36,37) and chemical exchange saturation transfer (CEST) imaging (38), and positron emission tomography (PET) with amino acid tracers, including [11C]-methyl-L-methionine (MET), (39) O-(2-[18F]-fluoroethyl)-L-tyrosine (FET) (40), 3,4-dihydroxy-6-[18F]-fluoro-L-phenylalanine (FDOPA) (41), also provide complementary diagnostic information in GBM detection. In addition, stimulated Raman scattering microscopy (42), optical coherence tomography (43), and mass spectroscopy (44) have also been developed to improve glioma diagnosis. However, most of these techniques do not have the capability of quantifying individual pathological components in a non-invasive manner.

To address limitations of conventional MRI, diffusion-weighted MRI-derived ADC has been one of the most widely researched tools for the evaluation of tumor cellularity and grade (10). Although increased tumor cellularity has been associated with decreased ADC, the expression of aquaporin in high-grade glioma (45), vasogenic edema (46), and necrosis (47) may obviate the interpretation of expected diffusion restriction caused by high cellularity. Indeed, in our tested tumor specimens, ADC did not correlate with cellularity while the DBSI-derived restricted fraction significantly correlated with both H&E and GFAP staining-based cellularity measures (Fig. 2). One observation in the present study contradicting the widely-accepted role of ADC in tumor cellularity is the significantly restricted diffusion observed in white matter tracts (Fig. 3), where the disrupted fiber network greatly increased diffusion restriction. Thus, our results further support that ADC alone cannot be considered a reliable tumor biomarker.

Through histological validation, we demonstrated DHI accurately detects and quantifies high tumor cellularity, tumor necrosis, and tumor infiltration. The newly-developed DHI framework accurately predicted key features of GBM microenvironment that eluded other neuroimaging technologies. Given the lack of specificity of clinical MRI in identifying tumor burden, DHI has the potential to aid in the non-invasive determination of tumor recurrence vs. treatment response. In addition, pre-operative DHI may help to guide biopsies and improve extent of resection.

## Acknowledgments

This work was supported in part by National Institutes of Health (R01-NS047592, P01-NS059560 and U01-EY025500 to S.-K.S., R01-NS094670 to A.H.K.), The Christopher Davidson and Knight Family Fund (A.H.K.), the Duesenberg Research Fund (A.H.K.), National Multiple Sclerosis Society (RG 1701-26617 to S.-K.S.), The Fundamental Research Funds for the Central Universities (SCUT 2018MS23 to R.Y.), Natural Science Foundation of Guangdong Province in China (2018A030313282 to R.Y.) and National Natural Science Foundation of China (81971574 to R.Y.).

## Author contributions

Z.Y., A.H.K., S.-K.S. and S.D. wrote the paper, and were assisted by R.L.P, J.L., R.H. and S.E.G. Z.Y., R.L.P. and D.D.M. performed the experiments. Z.Y., X.L., Q.Y., P.S., A.T.W., A.G., R.Y. and C.S. analyzed the experimental data. S.D. performed pathological evaluations. X.L., L.W. Z.Y., A.T.W. and J.-S.L. developed the classifier. M.W. conducted the statistical analyses. A.H.K. and R.P. provided surgical specimens for imaging. S.-K.S., A.H.K., S.D., J.-S.L. and J.L.C. oversaw the project. All authors have reviewed and approved the final version of the manuscript.

